# Seizures in a Dravet Syndrome mouse model inhibit the locus coeruleus but locally evoke supraphysiologic norepinephrine release

**DOI:** 10.64898/2026.09.25.754424

**Authors:** Meiling Zhao, Yiannos Demetriou, Anam Noor, Joseph Barden, Limei Zhu, Qianyi Yu, Alexander Sajdak, Arnav Akhaury, Chandni Rana, Peilin Yang, Yulong Li, Christian Burgess, Sam McKenzie, Paul Kramer, Joanna Mattis

## Abstract

Locus coeruleus (LC) neurons are the primary source of norepinephrine / noradrenaline (NE) released to most brain structures, including seizure-prone neocortical and limbic regions. LC-NE networks regulate numerous physiological processes, including arousal, attention, and memory. Previous evidence suggests that LC-NE neurons may be involved in epilepsy and that their activation may have therapeutic effects on seizures, but the endogenous activity of the LC-NE network during seizures remains largely unclear. We combined electrocorticography recordings and *in vivo* imaging to measure somatic activity of LC-NE neurons as well as downstream NE release during hyperthermia-induced seizures in a well-characterized pre-clinical model of Dravet Syndrome. We found that population LC-NE somatic activity was inhibited during seizures, whereas downstream NE release was dramatically elevated. To reconcile these discordant findings, we established that activation of downstream regions is capable of locally evoking NE release, in part via activation of glutamate receptors, consistent with the Glutamate Amplifies Noradrenergic Effects (GANE) model. Finally, we confirmed that ictal NE release occurred across additional downstream brain regions, epilepsy models, and seizure induction methods. Thus, disruption of NE signaling is likely a widespread ictal signature, with implications for impaired consciousness during and after seizures.

## Introduction

Impaired consciousness is a hallmark and highly burdensome feature of many epileptic seizures (Blumenfeld, 2012), often extending into the postictal period (Pottkämper et al., 2020). The ascending arousal system, a network of diffusely projecting brainstem neuromodulatory neurons, is an essential regulator of consciousness (French, 1957; Poe et al., 2020). Understanding the peri-ictal dynamics of these neuromodulatory circuits – including the Locus Coeruleus (LC) and its projections – is therefore essential for understanding how seizures disrupt consciousness, and for developing interventional therapies to preserve consciousness during and after seizures.

Multiple prior studies have investigated seizure-associated activity of LC noradrenergic (LC-NE) neurons (Silveira et al., 1998a, 1998b, 2000; Dutton et al., 2017), but data remain mixed as to whether LC-NE neurons are chiefly activated or inhibited, and primarily derive from low temporal resolution assays that limit interpretation. Several early studies employing chemical assays identified significant seizure-associated downstream release of noradrenaline / norepinephrine (NE) (Nelson et al., 1980; Sperk et al., 1983), but these also lacked the temporal resolution needed to distinguish between preictal, ictal, and postictal periods. A more recent study conducted *in vivo* single unit recordings in LC and fiber photometry recordings of a fluorescent NE reporter in hippocampus during perforant path stimulation-evoked seizures in rats (Larsen et al., 2023); they observed reduced ictal firing in the majority of LC-NE neurons, but a paradoxical large increase in fluorescent signal within the hippocampus.

The Glutamate Amplifies Noradrenergic Effects (GANE) model posits that glutamate spillover from excitatory synapses locally binds presynaptic glutamate receptors on NE axons, self-regulating NE release to selectively enhance the processing of high-priority information (Mather et al., 2016; Poe et al., 2020). We reasoned that – if substantiated – this model would provide a compelling candidate mechanism to explain ictal NE release absent significant LC activation.

Here, we sought to (1) record at high temporal resolution from LC-NE circuits in the well-characterized *Scn1a*-haploinsufficient mouse model of Dravet Syndrome (DS) (Mistry et al., 2014), and (2) directly test whether exogenous evoked activity within a region downstream of the LC is sufficient to elicit NE release *in vivo*.

By combining cortical electrocorticography (ECoG) with fiber photometry recordings, we found that hyperthermic seizures in DS mice triggered a population decrease in the somatic activity of LC-NE neurons, but an increase in NE sensor fluorescence within the hippocampal dentate gyrus (DG). We performed a series of control experiments to confirm that this downstream signal reflected NE release rather than a seizure-evoked optical artifact. We then demonstrated that sustained optogenetic stimulation within the DG triggered local NE release, absent somatic LC-NE activation, in part mediated by N-methyl-D-aspartate (NMDA) receptor activation, consistent with the GANE model. Finally, we demonstrated ictal release of NE across additional downstream regions, epilepsy models, and seizure induction methods. Taken together, our data support a model in which heightened ictal activity (i.e., within the seizure onset and propagation zones) directly acts upon local LC terminals, triggering supraphysiologic NE release.

## Methods

### Experimental animals

All animal procedures were approved by the University of Michigan Institutional Animal Care and Use Committee (IACUC). Mice were housed on a 12-hour light/dark cycle, with ad libitum access to food and water.

*Scn1a* haploinsufficient mice (*Scn1a^+/-^*; RRID: MMRRC_037107-JAX) were maintained on a 129S6.SvEvTac (129S) background. To generate experimental (DS) mice, 129S.*Scn1a^+/-^*males were crossed to (1) wild-type C57BL/6J females (B6; RRID:IMSR_JAX:033951), (2) females on a B6 background carrying the Dbh^tm3.2(cre)Pjen^/J transgenic allele (*Dbh-Cre*; RRID:IMSR_JAX:007914), or (3) *Dbh-Cre* females also carrying the Ai14 Cre-dependent tdTomato reporter allele (*Dbh-Cre*;*tdT*; RRID:IMSR_JAX:007914). Experimental progeny from each of these crosses had a 50:50 129S:B6 mixed background. *Scn1a^+/+^* (WT) littermates from respective crosses were used as experimental controls.

To generate focal cortical dysplasia (FCD) model mice, *in utero* electroporation was performed at embryonic day (E)14.5-15 on wild-type CD-1 background mice (RRID: IMSR_CRL:022), as previously described (Zhang et al., 2026). Mice were co-electroporated with (1) pCAG-RhebP37L (1.0 μg/μL), to introduce RHEBp.P37L, a patient-specific gain-of-function pathogenic mutation (Proietti Onori et al., 2021), and (2) pCAG-GFP (1.0 μg/μL), to fluorescently label the dysplasia.

Whole-cell patch-clamp recordings were performed in acute brain slices harvested from juvenile (postnatal day (P)21-P28) mice. All other experiments were performed in adult mice (at least P50). Male and female mice were used in equal proportion.

### Slice electrophysiology

#### Data acquisition

Whole-cell patch-clamp recordings were obtained from dopamine β-hydroxylase (DBH)-expressing locus coeruleus (LC) neurons as previously described (Zhu et al., 2025), with minor modifications. Mice were deeply anesthetized via isoflurane inhalation. Brains were rapidly removed and immersed in an ice-cold, carbogen-saturated (95% O_2_ / 5% CO_2_) sucrose-based cutting solution containing (in mM): 87 NaCl, 75 sucrose, 2.5 KCl, 1.0 CaCl_2_, 2.0 MgSO_4_, 26 NaHCO_3_, 1.25 NaH_2_PO_4_, and 10 glucose. Coronal brain slices (300 μm thick) containing the LC were prepared using a VT-1200S vibratome (Leica Microsystems). Slices were immediately transferred to a holding chamber filled with artificial cerebrospinal fluid (ACSF) containing (in mM): 125 NaCl, 2.5 KCl, 2.0 CaCl_2_, 1.0 MgSO_4_, 26 NaHCO_3_, 1.25 NaH_2_PO_4_, and 10 glucose, continuously bubbled with carbogen. Slices were allowed to recover at 32°C for 30 minutes, followed by an additional 30 minutes at room temperature (24°C) prior to electrophysiological recordings.

Slices were transferred to a recording chamber mounted on a SliceScope Pro 6000 electrophysiology system (Scientifica) and continuously perfused with oxygenated ACSF at a flow rate of ∼1 mL/min at 31 ± 1°C. Recording pipettes were pulled from borosilicate glass capillaries using a P-97 puller (Sutter Instruments) and had a tip resistance of 3–5 MΩ when filled with a K-gluconate-based internal solution. The internal solution consisted of (in mM): 130 K-gluconate, 6.3 KCl, 0.5 EGTA, 1.0 MgCl_2_, 10 HEPES, 4.0 Mg-ATP, and 0.3 Na-GTP (pH 7.30 adjusted with KOH, osmolarity 285 mOsm). Pipette positioning was controlled using a PatchStar micromanipulator (Scientifica). Series resistance was continuously monitored, and bridge balance was adjusted as necessary throughout the recording. No correction was made for the liquid junction potential.

Electrophysiologic signals were amplified using a MultiClamp 700B amplifier (Molecular Devices) and digitized via a Digidata 1550B interface (Molecular Devices) connected to a computer running Clampex 11.2 software. Signals were low-pass filtered at 10 kHz and sampled at 20 kHz. Resting potential measurements were recorded in current-clamp mode using a gap-free protocol without current injection. All other current-clamp recordings were obtained with the membrane potential adjusted to approximately −60 mV using a constant bias current. To evaluate individual action potential (AP) properties, a series of brief depolarizing current steps (10 ms duration, from +100 pA to +500 pA in 100-pA increments) were applied to elicit a single AP. To evaluate repetitive firing characteristics, a series of 1000-ms current injections ranging from −60 to +1200 pA in 20-pA increments was applied. Recordings with a series resistance exceeding 30 MΩ during the experiment were excluded from analysis.

#### Parameter extraction and statistical analysis

Analysis of electrophysiology data was performed using custom code: https://github.com/Joys-Dawn/EphysAutomatedAnalysis. Intrinsic membrane properties, AP parameters, and repetitive firing characteristics were calculated as previously described (Zhu et al., 2025). An AP was defined by the following criteria: (1) a threshold with a maximum derivative of voltage (dV/dt) > 10 mV/ms and (2) an amplitude crossing 0 mV.

Genotype comparisons of intrinsic electrophysiologic parameters were performed using linear mixed-effects models, with genotype included as a fixed effect and animal as a random intercept to account for correlations among cells obtained from the same animal. Degrees of freedom were estimated using the Satterthwaite approximation. P values were adjusted using the Benjamini-Hochberg false discovery rate procedure within each prespecified family of measurements (intrinsic membrane properties, individual AP properties, and repetitive firing properties).

### Stereotactic surgeries for virus injection and hardware implantation

Mice were anesthetized with 1-2% isoflurane throughout surgery and kept on a thermal blanket to maintain body temperature. Once deeply anesthetized (as determined by toe pinch), mice were secured to a stereotactic apparatus (Kopf Instruments). Craniotomies were made using an automatic drill (Kopf Instruments).

Adeno-associated viruses (AAVs) were employed to introduce molecular tools for manipulation and readout of circuit activity. To image LC-NE neuronal activity, Cre-dependent GCaMP8s (Zhang and Looger, 2024) was injected into LC (AAV9-syn-FLEX-jGCaMP8s-WPRE, 2.7 x 10^13^ genomic copies/mL, RRID:Addgene_162377). For the majority of NE imaging, the green NE sensor GRAB_NE2m_ (Feng et al., 2024) was injected into target regions (AAV9-hSyn-NE2m(NE3.1), 4.3 x 10^13^ genomic copies/mL, WZ Biosciences). A subset of experiments (as specified) included the red NE sensor (developed by Dr. Yulong Li’s laboratory; unpublished) (AAV2/9-hSyn-IgK-sw-rNE2.0, 1.01 x 10^13^ genomic copies/ mL, RRID:BrainVTA_PT-13850). Negative control mutant NE sensors included the green GRAB_NEmut_ (Feng et al., 2019) (AAV9-hSyn-NEmut(NE3.1mut), 2.0 x 10^13^ genomic copies/mL, WZ Biosciences) and the red GRAB_rNEmut_ (developed by Dr. Yulong Li’s laboratory; unpublished) (AAV2/9-hSyn-sw-rNEmut, 5.78 x 10^12^ genomic copies/ mL, BrainVTA). To evoke neuronal activity, the red-shifted excitatory opsin Chrimson (Klapoetke et al., 2014) was injected into LC (AAV5-syn-FLEX-rc[ChrimsonR-tdTomato], 2 x 10^13^ genomic copies/mL, RRID:Addgene_62723) or DG (AAV5-Syn-ChrimsonR-tdT, 1.2 x 10^13^ genomic copies/mL, RRID:Addgene_59171). As a negative control for photic stimulation, an mCherry-expressing virus was injected into DG (AAV5-hSyn-mCherry, 2.3 x 10^13^ genomic copies/mL, RRID:Addgene_114472).

All virus injections were performed using a 10 µL syringe and a 33-gauge beveled needle (Hamilton). Volumes were infused at a rate of 50 nL/minute, and the needle was left in place for 10 minutes before withdrawal. LC virus injection coordinates (relative to Bregma) were −5.2 mm (A/P), 1.0 (M/L), and −3.2 and −3.5 mm (D/V); 300 nL were infused at each depth. DG virus injection coordinates (relative to Bregma) were −2.0 mm (A/P), 1.3 mm (M/L), and −1.8 and −2.0 mm (D/V); 300 nL were infused at each depth. PVT virus injection coordinates (relative to Bregma) were −1.2 mm (A/P), 0.5 mm (M/L), and −2.95 mm (D/V), with a 10° angle towards the midline; 300 nL were infused. For injections in FCD mice, targeting was based upon gross visualization of electroporated fluorophore as seen under a fluorescent flashlight, with injections delivered immediately adjacent to the dysplastic region (cortex layer III/IV); 400 nL were infused. For fiber photometry and optogenetics experiments, a fiberoptic was then implanted on top of the virus injection site for fiber photometry recording (Thorlabs, 400 µm core diameter, 0.39 NA) and *in vivo* optogenetic stimulation (Doric, 200 µm core diameter, 0.22 NA).

For two-photon (2P) imaging in whisker (barrel) somatosensory cortex (wS1), a circular craniotomy (3.0 mm diameter) was made with the dura left intact. The center of the craniotomy was located over the wS1 cortex (3.5 mm lateral and 1.3 mm caudal relative to bregma). Virus was injected at three different locations on the cortical surface around the center of the craniotomy; 150 nL were injected per site. After the injection, the craniotomy was covered with a glass window made by gluing two pieces of coverslip glass (Harvard Apparatus) together. The smaller piece (3.0 mm diameter) was placed into the craniotomy while the larger piece (4.0 mm diameter) was glued to the bone surrounding the craniotomy.

For electrocorticography recordings, stainless steel skull screws (P1 Technologies) were implanted over the temporal and frontal cortices, with a reference electrode over the cerebellum. All mice used for fiber photometry and 2P recordings were additionally implanted with a head post (H.E. Parmer Co.) to enable head fixation. All implants were secured to the skull with a layer of dental cement (C&B Metabond) followed by dental acrylic (Fastray).

### Acute seizure induction

To evoke hyperthermic seizures, mice were head-fixed using a custom apparatus (Thorlabs) and placed under a heating lamp (Physitemp). Experimental animals were heated until the mouse exhibited a seizure or until the mouse internal body temperature reached 42.5°C, measured with a rectal probe (Physitemp). To reduce mortality, animals were then actively cooled (using external application of ice) until their internal body temperature recovered to at least 38 °C.

For chemoconvulsant seizures, pentylenetetrazole (PTZ) (MilliporeSigma, P6500) was dissolved in sterile saline to prepare a 100 mg/mL stock, and fresh aliquots were thawed and diluted with sterile saline to a working concentration of 10 mg/mL before each experiment. Mice were injected intraperitoneally with a dose of 60 mg/kg; if a seizure was not induced within approximately 5-10 minutes, a second dose of 30 mg/kg was administered.

### *In vivo* pharmacology

The α2-adrenergic receptor antagonist Yohimbine (Tocris, cat. no.1127) was reconstituted with 0.05% DMSO in saline and stored as a 0.5 mg/mL stock solution. The stock solution was thawed immediately before use and administered intraperitoneally at 2 mg/kg, 60 minutes before optogenetic stimulation.

The NMDA receptor antagonist 3-(2-carboxypiperazin-4-yl)propyl-1-phosphonic acid (CPP) (Tocris, cat. no. 0173) was reconstituted in sterile water and stored as a 5 mg/mL stock solution. The stock solution was thawed immediately before use and administered intraperitoneally at 10 mg/kg, 30 minutes before optogenetic stimulation.

### *In vivo* optogenetics

Yellow light was generated by a 574 nm laser (Shanghai Laser & Optics Century Co.) and delivered through a fiberoptic patch cord coupled to the implanted fiberoptic cannula via a connecting plastic sleeve (Precision fiber products). Laser output was controlled using a pulse generator (Master-9; A.M.P. Instruments).

### Electrocorticography (ECoG)

For simultaneous fiber photometry and ECoG recordings, ECoG signals were acquired and digitized with an Open Ephys data acquisition board (DAQ). For simultaneous 2P and ECoG recordings, ECoG signals were acquired and digitized with a National Instruments data acquisition card (ProAmp-8, CWE Inc.) and collected through Prairie View Imaging software from Bruker. All ECoG signals were sampled at a frequency of 1000 Hz.

Raw ECoG signal was bandpass filtered from 1 to 70 Hz using a fourth-order Butterworth filter, followed by a 60 Hz notch filter with a quality factor of 30. Filters were applied in both temporal directions to avoid phase displacement. Electrographic seizure onset was defined as ≥10 s of disrupted background with spike-wave discharges; offset was defined by cessation of discharges and the onset of consistent signal attenuation.

### Fiber photometry

#### Signal acquisition and synchronization

Fiber photometry signals were acquired at 100 Hz using a Fluorescence Mini Cube (Doric) connected to a PyPhotometry acquisition board. For all recordings from green sensors, LED excitation with 470 nm and 405 nm wavelengths was multiplexed to enable simultaneous acquisition of both signals. For simultaneous green and red sensor recordings, LED excitation was similarly multiplexed with 470 nm and 554 nm wavelengths.

When applicable, fiber photometry recordings were synchronized with ECoG recordings using shared TTL pulses. A linear fit between matched pulse times was used to correct clock drift, after which the 470 nm signal and 405 nm isosbestic control were interpolated onto the ECoG time base.

#### Preprocessing and z-scoring

For recordings with a 405 nm isosbestic control, both photometry channels were low-pass filtered at 3 Hz. The filtered isosbestic signal was fitted to the filtered 470 nm signal by iteratively reweighted least-squares (IRLS) regression (Keevers and Jean-Richard-dit-Bressel, 2025). Regression weights were calculated using Tukey’s bisquare function with a tuning constant of 1.4.

The fitted isosbestic signal was calculated as

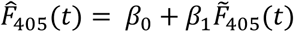

and fractional fluorescence change was calculated as

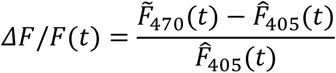

For simultaneous green and red sensor recordings, each channel was processed independently using a fourth-order Butterworth low-pass filter with a 10 Hz cutoff, applied in both temporal directions. A biexponential decay function was fitted to each filtered fluorescence trace to estimate the photobleaching baseline. For each channel, ΔF/F was calculated by subtracting the fitted baseline from the filtered fluorescence signal and dividing by the fitted baseline.

The resulting ΔF/F trace was standardized via z-scoring within each recording using the mean and standard deviation from the beginning of the analyzed recording to the onset of heating.

PTZ recordings were z-scored using the mean and standard deviation of the interval from recording onset to PTZ administration (120 seconds). For optogenetic recordings the normalization period was the first 100 seconds of the recording.

Recordings comparing evoked signal before versus after drug (Yohimbine and CPP) administration were not z-scored because pharmacologic manipulations may impact the baseline as well as evoked response (Wallace et al., 2025).

Analysis of ECoG and fiber photometry data was performed using custom code: https://github.com/Joys-Dawn/Fiber-Photometry-ECoG.

#### Visualization and quantification of seizure and opto-evoked responses

For hyperthermia recordings that evoked seizures, the baseline (interictal) period was defined from recording onset to heating onset. The preictal period was defined as the final 10 seconds preceding seizure onset. The ictal period was defined from electrographic seizure onset to offset.

For hyperthermia recordings that failed to evoke seizures, equivalent event times were assigned using the associated seizure cohort. Equivalent seizure onset times were defined as the first points during heating at which the animal’s temperature was closest to the cohort mean onset temperature. Equivalent offset was defined as equivalent onset plus the mean seizure duration.

For ictal recordings (and their associated controls), event-aligned photometry traces were extracted around seizure onset (or equivalent temperature). Traces were baseline-subtracted using the mean signal from 5 to 1 seconds before each event. Area under the curve (AUC) was calculated by trapezoidal integration from seizure or event onset. For LC GCaMP recordings, AUC was calculated over 100 seconds after seizure (or equivalent) onset. For DG GRAB_NE2m_ recordings, the integration window matched the duration of each seizure, including positive signal values only; non-seizure recordings used the mean seizure duration. Ictal maxima and minima were defined as the highest and lowest signal levels sustained for at least 200 ms. Changes in these measures were calculated relative to the mean standardized ΔF/F during the 10 seconds preceding seizure onset.

For optogenetic experiments involving a single burst of photic stimulation, event-aligned photometry traces were extracted around the stimulation onset. The mean signal from 5 to 1 seconds before stimulation was subtracted from each trace, and the maximum, minimum, and signed area under the curve were calculated over the 40 seconds following stimulation onset.

For optogenetics experiments involving repeated bursts of photic stimulation, the mean ΔF/F during the full period preceding the first train was subtracted from each recording. Event-aligned traces were extracted from 5 seconds before to 10 seconds after each train onset, and the maximum ΔF/F and signed area under the curve were calculated over the 10 seconds following onset. Train-level measurements were averaged to obtain one value per mouse and condition.

#### Transient detection and parameter extraction

Baseline LC-NE GCaMP recordings were further analyzed to characterize calcium transients. Transients were detected using a two-stage procedure adapted from the Prominence Method (Wallace et al., 2025). Candidate peaks were first identified in the standardized ΔF/F trace using a minimum height of 1 SD. Consecutive peaks were required to be separated by at least 0.3 seconds. Each candidate was then evaluated according to its prominence in the fractional ΔF/F trace, with minimum prominence set at 0.015 ΔF/F.

Peaks with a half-width greater than 8 s were excluded. Transient amplitude was defined as the difference between the signal at the peak and the minimum signal within 2.5 seconds on either side of the peak. Transient half-width was measured at 50% prominence in the standardized ΔF/F trace. Transient frequency was calculated as the number of detected events divided by the duration of the analysis interval.

### Statistical analysis

Each recording session contributed one value for each scalar outcome. Between-group comparisons evaluated: (1) DS versus WT, for baseline transient measures in the LC GCaMP dataset; (2) seizure versus no seizure, for the primary LC GCaMP and DG GRAB_NE2m_ outcomes; and (3) pre-versus post-drug, for the Yohimbine and CPP experiments.

The distribution of each outcome was assessed separately within each group. Non-parametric tests were used when any group had an absolute skewness of 2 or more, an absolute excess kurtosis of 7 or more, or a Shapiro-Wilk p-value of less than 0.05. Two-group comparisons were performed using two-sided Welch unequal-variance t-tests when the parametric criteria were met. Two-sided Mann-Whitney U tests were used otherwise. Within-session comparisons of heating and ictal values were performed using paired t-tests when the paired differences met the distributional criteria and Wilcoxon signed-rank tests otherwise.

Group summaries are presented as mean ± standard error of the mean (SEM). Statistical significance was evaluated at alpha=0.05. Analyses were performed using custom Python code with NumPy, SciPy, and pandas.

### Acquisition and analysis of 2P data

*In vivo* 2P imaging was performed in whisker (barrel) somatosensory cortex (wS1) on head-fixed mice. Images were acquired on a Bruker 2P microscope equipped with an 8 kHz resonant scanning module, a 16 × 0.8 NA microscope objective (Nikon). GRAB_NE2m_ was excited at 960 nm (40-60 mW at specimen) with an InSight X3 tunable ultrafast Ti:Sapphire laser (Spectra-Physics). Imaging fields were restricted to areas where GRAB_NE2m_ expression overlapped with the center of the cranial window (3.5 mm lateral and 1.3 mm caudal to bregma). The beam was focused on 150-250 μm from the cortical surface. Images were acquired with a resolution of 512 × 512 pixels at 30 Hz using Prairie View Imaging.

Images were processed using the Suite2P pipeline in Python (Stringer et al., 2019). After correcting for motion, regions of interest (ROIs) were manually selected. A double exponential fit was used to remove the influence of photobleaching. The signal was then z-scored across the 80 seconds preceding seizure onset.

### Acquisition and analysis of live imaging in acute slice

#### Tissue sectioning

Mice were anesthetized using isoflurane, decapitated, and brains were rapidly extracted. Coronal sections were cut at 330 µm thickness on a vibratome (Campden Instruments) while immersed in ice-cold slicing solution containing (in mM): 198 glycerol, 2.5 KCl, 1.2 NaH_2_PO_4_, 25 NaHCO_3_, 20 HEPES, 10 glucose, 10 MgCl_2_ and 0.5 CaCl_2_. Sectioned tissues were immediately removed from the slicing ACSF and moved to a holding chamber in a heated water bath (34°C). Sections were incubated for 30 minutes in a holding solution containing (in mM): 92 NaCl, 2.5 KCl, 1.2 NaH_2_PO_4_, 30 NaHCO_3_, 20 HEPES, 35 glucose, 2 MgCl_2_, 2 CaCl_2_, 5 sodium ascorbate, 3 sodium pyruvate and 2 thiourea. Throughout the entire process, 95% O_2_ and 5% CO_2_ were saturated through the slicing ACSF as well as the holding ACSF. Following incubation, the holding chamber was placed at room temperature and used for experiments for up to 8 hours. During experiments, slices were transferred to a heated (33°C - 35°C) recording chamber which was continuously perfused with ACSF containing (in mM): 125 NaCl, 25 NaHCO_3_, 1.25 NaH_2_PO_4_, 3.5 KCl, 10 glucose, 1 MgCl_2_ and 2 CaCl_2_, saturated with 95% O_2_ and 5% CO_2_.

#### Live imaging

Brain sections were imaged on an upright microscope (Scientifica, UK) with a 10x, 0.25 NA Plan Achromat objective (Evident, Japan). Brain tissue was illuminated with a 450 nm LED (CoolLED, UK), and emitted light was isolated with a GFP filter cube set (Thorlabs, USA). An sCMOS camera (Prime BSI Express, Teledyne, Canada) was used to visualize the tissue, and images were captured on a computer using Micro-Manager software (Edelstein et al., 2014). Images were measured using a 1 mm stage micrometer to calibrate the objective.

#### Image analysis and statistics

Images were analyzed using FIJI (Schindelin, J., 2012). Images were registered using the Linear Stack Alignment with SIFT plug-in (Lowe, D.G., 2004). An ROI was drawn around the DG and the average pixel intensity was measured for each condition. Each section was then normalized to its own baseline to account for differences in GRAB_NE2m_ expression levels between sections and animals. Statistics were performed using GraphPad Prism version 10.4.1 for Windows (GraphPad Software, Boston, Massachusetts, USA; www.graphpad.com). Results were reported as mean ± SEM.

### Immunohistochemistry and fluorescent imaging of fixed tissue

For histological confirmation of expression of viral constructs and immunohistochemical labeling of tyrosine hydroxylase (TH), brains were harvested immediately after death (resulting either from SUDEP or isoflurane overdose followed by transcardial perfusion with ice-cold PBS then 4% paraformaldehyde (PFA)). Tissue was post-fixed overnight in 4% PFA, then transferred to 30% sucrose in PBS and allowed to equilibrate for approximately 24 hours. 40 μm sections were obtained using a sliding microtome (Leica Biosystems model SM2010 R) and stored in a cryoprotectant solution (25% glycerol and 30% ethylene glycol in PBS, pH 6.7) at −20 °C.

Free-floating sections encompassing LC were stained for TH as follows: Sections were washed three times in PBS then incubated in a blocking solution of 3% normal donkey serum (RRID:AB_2337258, Jackson ImmunoResearch, cat. no. 005-000-121) and 0.3% Triton™ X-100 (MilliporeSigma cat. no. X100-100ML) for 30 minutes. Sections were then incubated overnight at 4 °C in a rabbit polyclonal anti-TH antibody (1:1000; RRID: AB_390204, MilliporeSigma, cat. no. AB152). This was followed by three additional washes in PBS, then a 3-hour incubation with an anti-rabbit secondary antibody (1:500, RRID:AB_2492288, Alexa Fluor® 647 AffiniPure Donkey Anti-Rabbit; Jackson ImmunoResearch, cat. no. 711-605-152). Sections were washed three more times in PBS, then mounted using a medium with DAPI (Vectashield® PLUS Antifade Mounting Medium with DAPI; Vector Laboratories cat. no. H2000) or without (Polyvinyl Alcohol Mounting Medium with DABCO™, Antifading pH 8.7; MilliporeSigma Supelco cat. no. 10981) as appropriate. All images were obtained using a Nikon fluorescent microscope and NIS-Elements software.

To assess activation of TH-expressing neurons in the LC following seizure, adult DS mice underwent hyperthermia to induce seizure alongside WT littermates for comparison. Approximately 90 minutes following the hyperthermia procedure, mice were sacrificed and transcardially perfused and brain tissue was harvested as described above. Then, free-floating sections containing LC were stained for c-Fos and counterstained for TH using the same procedure described above, but with the following primary antibodies: rabbit monoclonal anti-c-Fos (1:1000; RRID:AB_2247211, Cell Signaling Technology, cat. no. 2250) and mouse monoclonal anti-TH (1:500; Invitrogen, cat. no. MA1-24654). For visualization, secondary antibodies against rabbit (1:500; RRID:AB_2313584, Alexa Fluor® 488 AffiniPure Donkey Anti-Rabbit, Jackson ImmunoResearch, cat. no. 711-545-152) and mouse (1:500; RRID:AB_2340855, Alexa Fluor® 594 AffiniPure Donkey Anti-Mouse, Jackson ImmunoResearch, cat. no. 715-585-151) were used. To quantify the proportion of c-Fos-positive cells within the LC, z-stacks were captured using a Nikon fluorescent microscope and cells expressing either c-Fos, TH, or both were manually counted in NIS-Elements.

## Results

### Population LC-NE somatic activity is inhibited by seizures

To determine the peri-ictal dynamics of LC-NE neurons, we selectively injected Cre-dependent GCaMP8s (Zhang and Looger, 2024) into the LC of *Scn1a*^+/-^;*Dbh-Cre* (DS) mice and *Scn1a*^+/+^;*Dbh-Cre* (WT) littermate controls and performed fiber photometry recordings concurrently with cortical ECoG (**Fig. 1A-B; Supplemental Fig. 1**).

**Figure 1.**
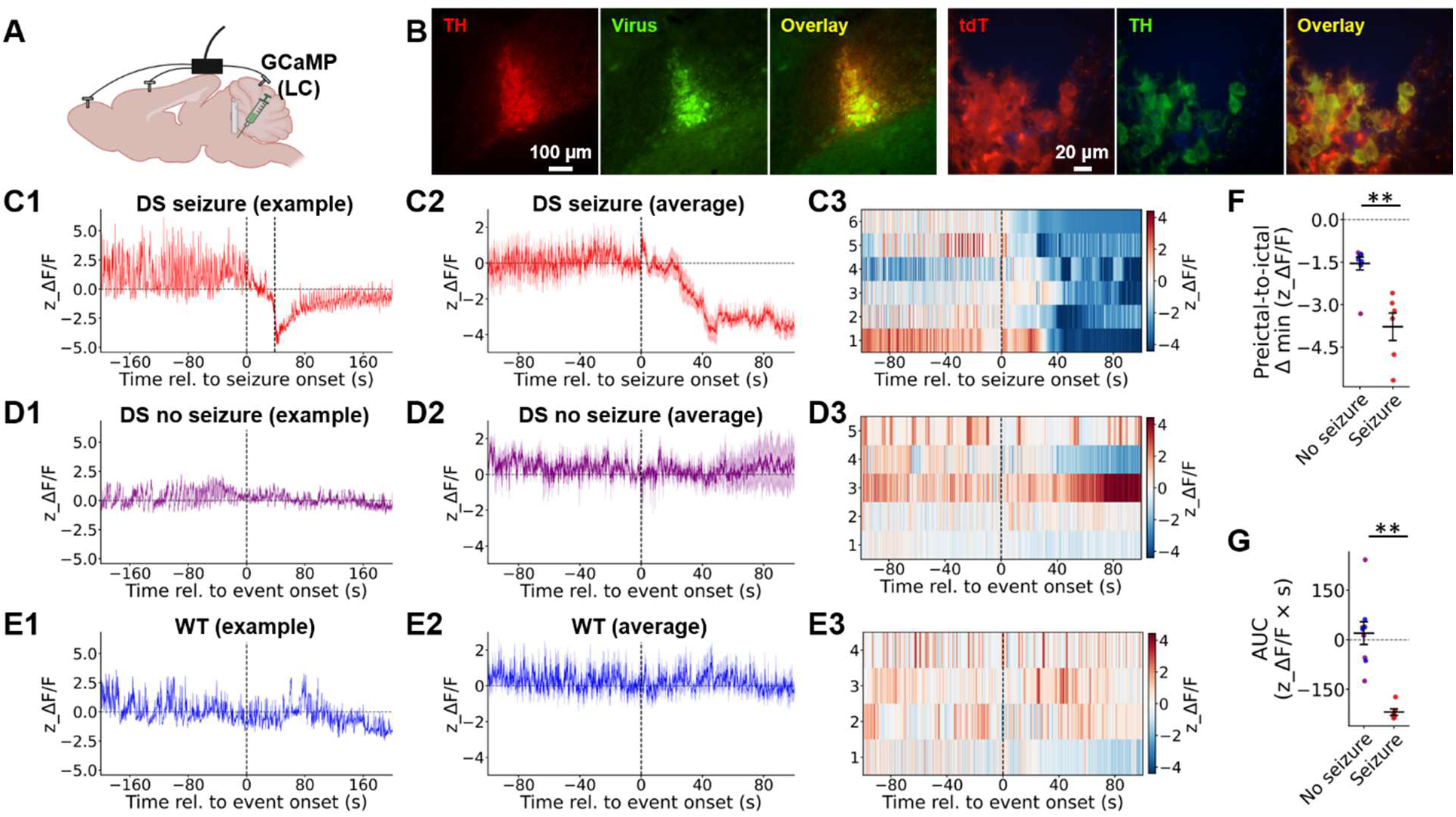
Seizures reduce population LC-NE somatic activity. **(A)** Schematic illustrating experimental configuration for LC-NE somatic recordings, including cre-dependent-GCaMP8s virus injection into LC, fiberoptic placement above LC, and cortical skull screws for ECoG. **(B)** Left: low magnification images illustrating cre-dependent viral expression (green) within the LC, as delineated by TH immunostaining (red). Right: high magnification images illustrating high specificity of the *Dbh-Cre* line, based on colocalization of tdT reporter expression (red) with TH immunostaining (green) (119 / 122 colocalized cells (∼98%) across n = 7 sections / 3 mice.) **(C)** Fiber photometry GCaMP recordings from LC-NE neurons during sessions in which DS mice exhibited seizures (n = 5 mice / 6 sessions). **(D)** Recordings from DS mice that exhibited no seizures (n = 4 mice / 5 sessions). **(E)** Recordings from WT mice (n = 3 mice / 4 sessions). Displayed are individual traces **(C1, D1, E1)**, mean (dark lines) and SEM (shaded areas) across all sessions within the group **(C2, D2, E2)**, and heatmaps of z_ΔF/F values, in which each row represents data from an individual session **(C3, D3, E3)**. Traces are centered on either seizure onset **(C)** or event onset defined by an equivalent mean temperature **(D, E)**; the second vertical line in **C1** indicates seizure offset in that individual recording. **(F)** Change in z_ΔF/F in seizure sessions (−3.78 ± 0.48) versus sessions without seizures (−1.55 ± 0.23; p = 0.003, Mann-Whitney U test). **(G)** AUC calculated across the 100 s after seizure onset from the z_ΔF/F plots. The AUC was significantly smaller (more negative) in seizure sessions (−218.80± 9.68) than in sessions without seizures (20.04 ± 34.40; p = 0.0004, Mann-Whitney U test). Note that for **F-G**, data were pooled across both DS (purple) and WT (blue) sessions that failed to evoke seizures.

Interictally (i.e., at room temperature), we observed no genotype differences in spontaneous calcium transients (**Supplemental Fig. 2A-B**). Consistent with this, whole-cell patch-clamp recordings from LC-NE neurons revealed no genotype differences in intrinsic properties or action potential firing (**Supplemental Fig. 2C-D; Supplemental Table 1**).

Naturalistic bilateral tonic-clonic seizures can be readily evoked in DS mice by hyperthermia, phenocopying the febrile seizures that prominently define the clinical condition (Millichap et al., 2009). We heated DS mice until electroclinical seizure onset and observed a significant subsequent decrease in the LC-NE GCaMP signal (**Fig. 1C**). No such decrease was evident in the isosbestic channel (**Supplemental Fig. 1A**), arguing against a calcium-independent artifact. A subset of hyperthermia sessions in DS mice failed to trigger seizures prior to the experimental endpoint (42.5 °C internal body temperature); these sessions, as well as recordings in WT mice, served as negative controls for the effect of heating alone, and did not demonstrate equivalent changes (**Fig. 1D-E; Supplemental Fig. 1B**).

To further distinguish seizure-associated changes from heating-related changes at comparable temperatures, we aligned the GCaMP signal either to seizure onset when present, or, in sessions without seizures, to the temperature equivalent to the mean seizure onset threshold (here termed “event onset”). To quantify the seizure-evoked signal decrease, we calculated (1) the difference between the pre-ictal mean and the ictal minimum (**Fig. 1F**), and (2) the area under the curve (AUC) of the fluorescence (z_ΔF/F) across the first 100 s following seizure onset (**Fig. 1G**). For “no seizure” DS and WT sessions, we performed the same calculations, using the matched, temperature-equivalent time values. Together, these results demonstrated a significant seizure-associated decrease in LC-NE neuron activity.

### Seizures evoke supraphysiologic NE release in the DG

We next investigated the peri-ictal dynamics of downstream NE release within this same experimental paradigm. We expressed the fluorescent GPCR-Activation-Based (GRAB) sensor, GRAB_NE2m_ (Feng et al., 2024) within the dorsal dentate gyrus (DG), a major downstream target of LC projections (Blackstad et al., 1967), and again performed simultaneous fiber photometry and ECoG recordings while evoking seizures with hyperthermia (**Fig. 2A-B**).

**Figure 2.**
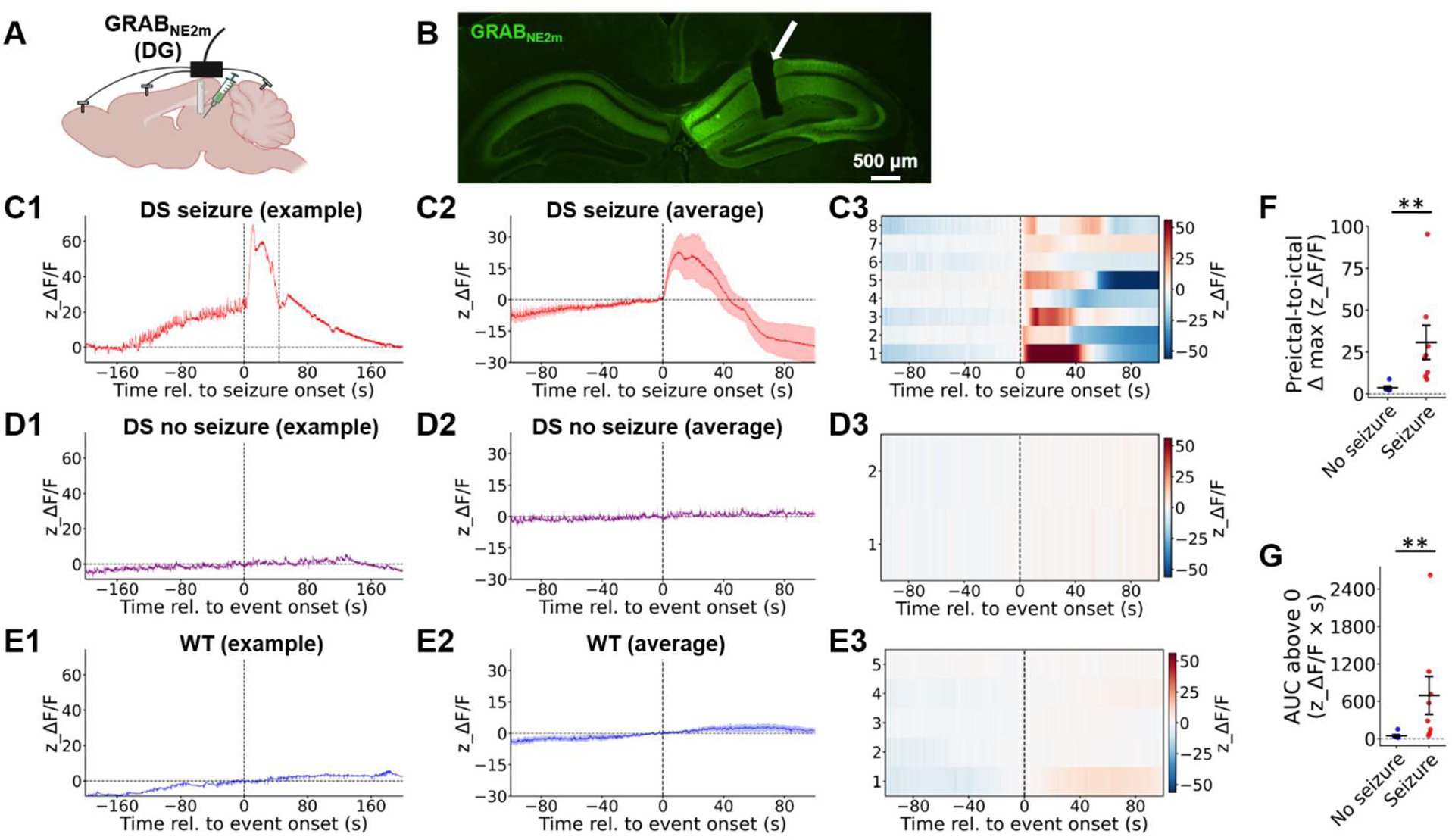
Seizures trigger a large release of NE within DG. **(A)** Schematic illustrating experimental configuration for NE recordings in DG, including GRAB_NE2m_ virus injection into DG, fiberoptic placement above DG, and cortical skull screws for ECoG. **(B)** Expression of GRAB_NE2m_ (green) virus within DG. Note the fiberoptic track (arrow). **(C)** Fiber photometry GRAB_NE2m_ recordings from DG during sessions in which DS mice exhibited seizures (n = 7 mice / 8 sessions). **(D)** Recordings from DS mice that exhibited no seizures (n = 2 mice / 2 sessions). **(E)** Recordings from WT mice (n = 3 mice / 5 sessions). Displayed are individual traces **(C1, D1, E1)**, mean (dark lines) and SEM (shaded areas) across all sessions within the group **(C2, D2, E2)**, and heatmaps of heatmaps of z_ΔF/F values, in which each row represents data from an individual session **(C3, D3, E3)**. Traces are centered on either seizure onset **(C)** or event onset defined by an equivalent mean temperature **(D, E)**; the second vertical line in **C1** indicates seizure offset in that individual recording. **(F)** Change in z_ΔF/F value from the immediate pre-ictal (or equivalent) time period to the maximum value within the ictal (or equivalent) time period. The signal increase was significantly greater in seizure sessions (30.73 ± 10.17) than in sessions without seizures (3.67± 0.87; p = 0.0006, Mann-Whitney U test). **(G)** AUC calculated across the positive section after seizure onset for each individual session from the z_ΔF/F plots. The AUC ranged from 50.23 to 2622.40 in seizure sessions (mean 694.16 ± 302.68) and from 15.41 to 150.24 in sessions without seizures (mean 45.19 ± 17.80; p = 0.002, Mann-Whitney U test). Note that for **F-G**, data were pooled across both DS (purple) and WT (blue) sessions that failed to evoke seizures.

Seizures in DS mice were accompanied by a rapid and dramatic increase in GRAB_NE2m_ signal (**Fig. 2C**). When z-scored relative to the baseline (pre-heating) signal, the mean magnitude of this ictal increase was 18.1 (range 2.3-95.4) – i.e., dramatically outside of the typical physiologic domain. No such signal changes were observed in either DS or WT heating sessions without seizures (**Fig. 2D-G**).

Given the discordance between reduced somatic LC-NE neuronal activity (**Fig. 1**) and increased downstream NE release (**Fig. 2**), we conducted multiple control experiments to confirm that the GRAB_NE2m_ signal was indeed encoding NE. In WT mice, we expressed GRAB_NE2m_ within DG, and the red-shifted excitatory opsin, Chrimson, within LC-NE neurons, to enable photoactivation within LC and fiberoptic recordings within DG (**Fig. 3A-B**). Optogenetic activation of LC-NE neurons reliably and repeatedly triggered robust GRAB_NE2m_ signal in DG, which was largely abolished following systemic injection of yohimbine, an α2-adrenergic receptor antagonist that competitively prevents NE binding to GRAB_NE2m_ (Feng et al., 2019) (**Fig. 3C**). Quantification of responses across the first 10 light pulses with versus without Yohimbine showed significant decreases in peak evoked amplitudes (0.14 ± 0.03 pre versus 0.03 ± 0.01 post; p = 0.013; paired t-test) and AUC (1.03 ± 0.25 pre versus 0.15 ± 0.04 post; p = 0.016; paired t-test; n = 5 mice / 5 paired sessions). For further tool validation, we performed live imaging in GRAB_NE2m_-expressing acute slices, confirming that wash-on of NE (50 µM) evoked a large increase in fluorescence in slices pre-treated with vehicle (DMSO; 0.2% in ACSF), but not in slices pre-treated with Yohimbine (20 µM) (**Fig. 3D**).

**Figure 3.**
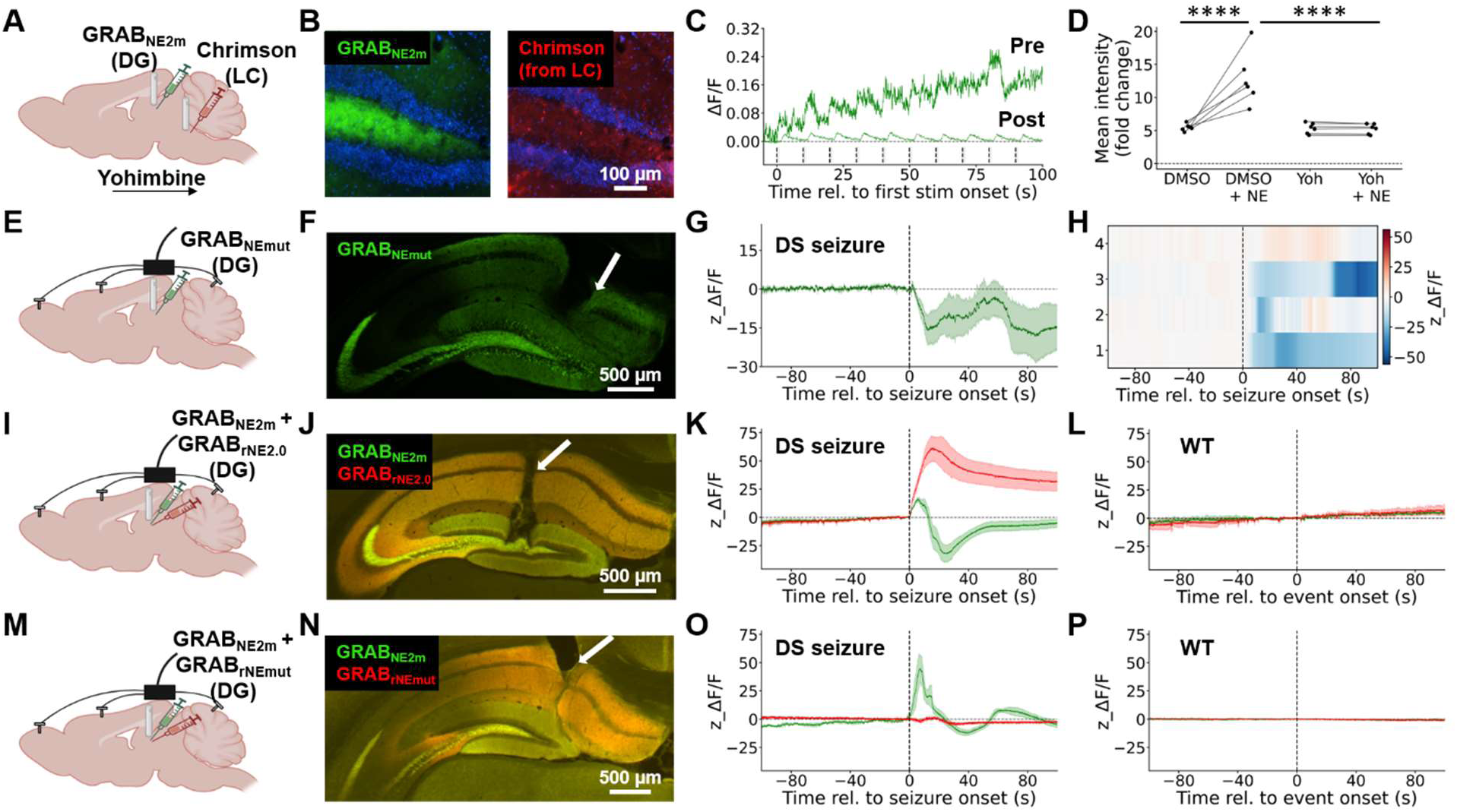
Validation of the GRABNE2m sensor during ictal recordings. **(A)** Schematic illustrating experimental configuration for concurrent optogenetic activation and fiber photometry readout of the LC-NE circuit, including cre-dependent Chrimson injection into LC, GRAB_NE2m_ virus injection into DG, and fiberoptic placement above both LC (for photic stimulation) and DG (for fiber photometry). **(B)** Confirmation of GRAB_NE2m_ expression (green) and LC fibers (red) within the DG. **(C)** Representative GRAB_NE2m_ responses recorded in DG in response to repeated photic stimulation in LC (10 mW, 5 ms pulse width, 20 Hz), pre (upper trace) versus post (lower trace) systemic Yohimbine injection. **(D)** Quantified fluorescence intensity within live-imaged GRAB_NE2m_-expressing acute slices after wash-on of (1) DMSO or Yohimbine followed by (2) DMSO + NE or Yohimbine + NE. There was a significant interaction between initial drug and pre versus post NE fluorescence levels (F(1,10)=20.11, p=0.0012, repeated measures 2-way ANOVA). Stars (****) indicate post-hoc pair-wise tests with significance of p < 0.0001 (Fisher’s Least Significant Difference (LSD) test). **(E)** Schematic illustrating experimental configuration for NE mutant virus recordings in DG, including GRAB_NEmut_ virus injection into DG, fiberoptic placement above DG, and cortical skull screws for ECoG. **(F)** Representative expression of GRAB_NEmut_ within DG; note the fiberoptic track (arrow). **(G)** Mean (dark lines) and SEM (shaded areas) of population GRAB_NEmut_ signal (n = 3 mice / 4 sessions). **(H)** Heatmap of GRAB_NEmut_ z_ΔF/F values, in which each row represents data from an individual session. No GRAB_NEmut_ recordings showed a seizure-evoked signal increase. **(I)** Schematic illustrating experimental configuration for dual NE sensor recordings in DG, including dual GRAB_NE2m_ and GRAB_rNE2.0_ virus injection into DG, fiberoptic placement above DG, and cortical skull screws for ECoG. **(J)** Co-expression of GRAB_NE2m_ and GRAB_rNE2.0_ in the DG; note the fiberoptic track (arrow). **(K)** Both GRAB_NE2m_ (green) and GRAB_rNE2.0_ (red) signal showed an elevation following seizure onset (n = 2 mice / 4 sessions). **(L)** Neither GRAB_NE2m_ nor GRAB_rNE2.0_ signal changed in heated WT mice (n = 2 mice / 2 sessions). **(M)** Schematic illustrating experimental configuration for dual GRAB_NE2m_ and GRAB_rNEmut_ virus injection into DG, fiberoptic placement above DG, and cortical skull screws for ECoG. **(N)** Co-expression of GRAB_NE2m_ and GRAB_rNEmut_ in the DG; note the fiberoptic track (arrow). **(O)** GRAB_NE2m_ (green) signal increased after seizures in DS mice, but GRAB_rNEmut_ (red) did not change (n = 2 mice / 4 sessions). **(P)** Neither GRAB_NE2m_ nor GRAB_rNEmut_ signal changed in heated WT mice without seizures (n = 2 mice / 3 sessions).

We next performed additional control experiments designed specifically to validate the GRAB_NE2m_ signal within the ictal context. First, we recorded during seizures from a mutant NE sensor (GRAB_NEmut_) (Feng et al., 2019), which harbors a point mutation to prevent NE binding. We observed no seizure-associated signal increase in this mutant sensor, an important negative control result supporting that the ictal GRAB_NE2m_ fluorescence peak (**Fig. 2**) reflects true NE release rather than a seizure-associated artifact. However, the mutant sensor did demonstrate a significant, transient reduction in fluorescence following seizure onset (**Fig. 3E-H**). To understand this response, we scrutinized both the raw 470 nm and 405 nm (isosbestic) signals. The GRAB_NE2m_ sensor showed a large seizure-associated increase in the 470 nm signal (**Supplemental Fig. 3A**) – underlying the positive signal in the corrected trace, as described (**Fig. 2**) – but in fact a transient decrease in the 405 nm signal. In contrast, the GRAB_NEmut_ sensor exhibited a seizure-associated decrease in both signals, consistent with artifact (**Supplemental Fig. 3B**).

Having identified this ictal artifact in the GRAB_NE2m_ sensor, we opted to additionally measure peri-ictal NE release with the newly developed, red-shifted NE sensor, GRAB_rNE2.0_. We co-expressed GRAB_NE2m_ and GRAB_rNE2.0_ within the DG and recorded both channels simultaneously during hyperthermia sessions in DS and WT mice (**Fig. 3I-L**). As GRAB_rNE2.0_ has no reported isosbestic wavelength, we co-injected (in a separate cohort) GRAB_NE2m_ and the red mutant (NE-insensitive) sensor, GRAB_rNEmut_ (**Fig. 3M-P**). The GRAB_rNE2.0_ sensor exhibited a dramatic signal increase following seizure onset (**Fig. 3K**) that was absent across all negative controls. The ictal increase in signal was even more dramatic with GRAB_rNE2.0_ than with GRAB_NE2m_; while the GRAB_NE2m_ response was biphasic (an initial increase, followed by a decrease in fluorescence), the GRAB_rNE2.0_ response showed only the initial increase.

Collectively, these results demonstrate that the negative deflection in the GRAB_NE2m_ signal derives from an artifactual (i.e., calcium-independent) change in sensor function within the ictal context. This negative deflection reduces the peak signal magnitude and complicates detailed interpretation of ictal NE dynamics. However, these control experiments also support the central conclusion that the positive component of the fluorescent GRAB signal reflects a true and significant increase in ictal NE release.

### NE release is locally evoked by downstream neuronal activity, and partially mediated by NMDA receptors

The discordance between LC-NE somatic activity and DG NE release could potentially arise from alternative noradrenergic projections to DG. We considered this explanation unlikely, as the LC is thought to be the sole source of DG NE (Szabadi, 2013). The nucleus tractus solitarius (NTS) contains NE neurons that functionally project to the amygdala (Clayton and Williams, 2000) but we found no evidence of projections between the NTS and DG (**Supplemental Fig. 4**).

A second potential explanation for the discordant ictal NE release is that a subset of LC-NE neurons are highly activated by seizures, but that this activation is obfuscated by the bulk fiber photometry signal. We quantified expression of the immediate early gene, c-Fos, within LC-NE neurons following either evoked seizures (in DS mice) or equivalent hyperthermia exposure (WT controls) and found no difference between groups (**Supplemental Fig. 5**).

Alternatively, activity within DG (as occurs during seizures in DS mice (Liautard et al., 2013)) may directly trigger NE release. To confirm that NE release can be locally evoked, we co-expressed GRAB_NE2m_ and Chrimson within the DG (**Fig. 4A**), and measured NE release in response to photic stimulation. We found that 10 seconds of 20 Hz DG photostimulation evoked profound NE release (**Fig. 4B**). Within the same animals, 1 Hz stimulation evoked no response (**Fig. 4C**), indicating that this process was dose or frequency-dependent, and may be less likely to occur under physiological regimes. To ensure that the results were not attributable to inadvertent recruitment of LC-NE somatic activity, we additionally expressed GCaMP8s within LC-NE neurons in a subset of the same animals. No significant Chrimson expression was histologically detected within the LC (**Fig. 4D**), and photoactivation of DG Chrimson had no appreciable effect on LC GCaMP signal (**Fig. 4E**), arguing against either direct or polysynaptic recruitment of LC-NE cells under the same conditions that robustly evoked NE release. Finally, as a negative control for direct light effects, we confirmed absence of evoked GRAB_NE2m_ signal with 20 Hz DG photostimulation in mice expressing a fluorophore control virus (**Fig. 4F-G**).

**Figure 4.**
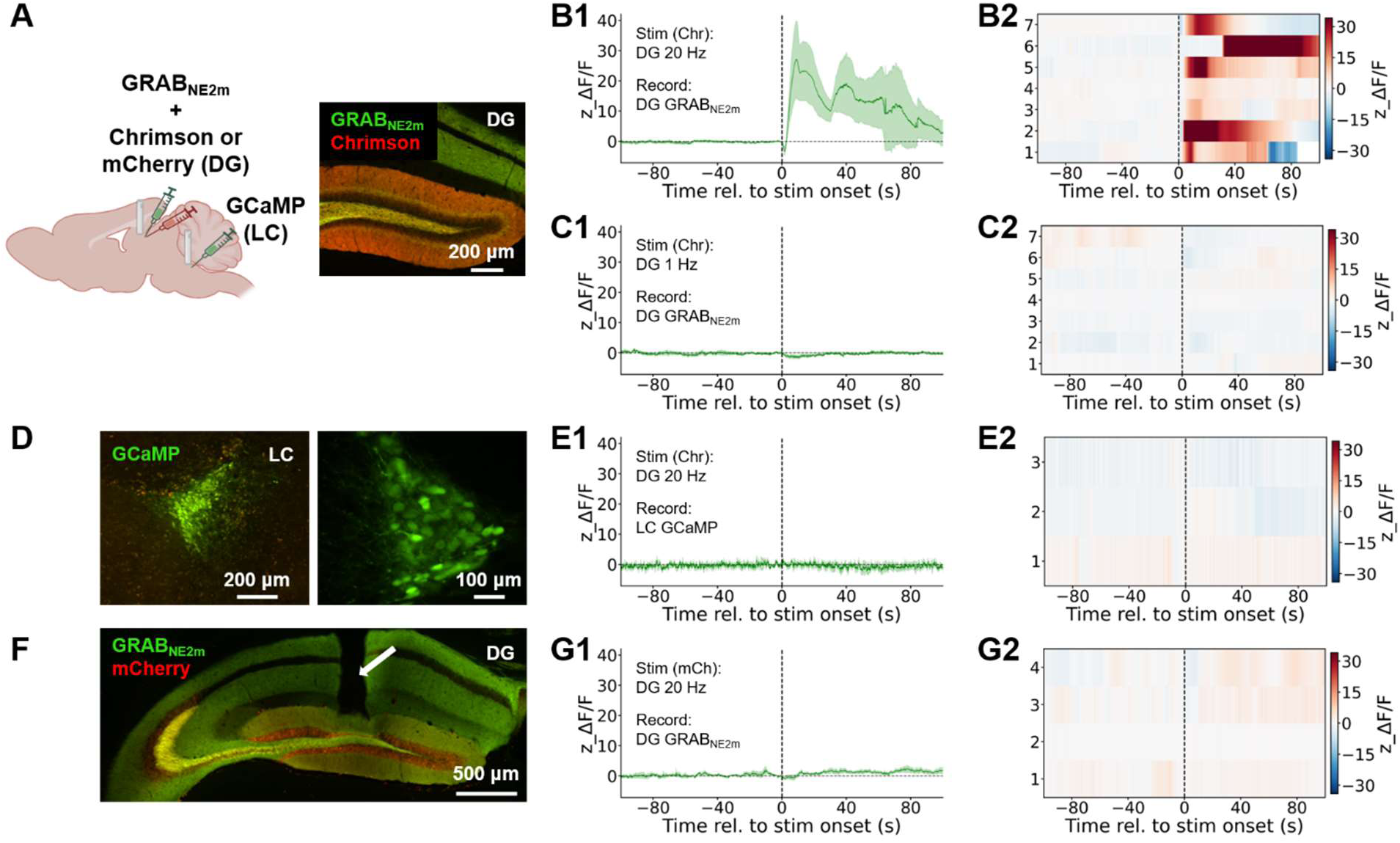
Local activity within the DG evokes NE release independent of LC somatic activation. **(A)** Left: Schematic illustrating experimental configuration for local (DG) optogenetic stimulation and concurrent LC-NE circuit recordings, including combined expression of GRAB_NE2m_ and either Chrimson or mCherry in DG, expression of cre-dependent GCaMP in LC, and fiberoptic placement above both DG (for photic stimulation and fiber photometry) and LC (for fiber photometry). Right: Co-expression of GRAB_NE2m_ (green) and Chrimson (red) in the DG. **(B)** Robust DG GRAB_NE2m_ response to 20 Hz photic activation of Chrimson within DG (n = 7 mice / 7 sessions). **(C)** Lack of DG GRAB_NE2m_ response to 1 Hz photic activation of Chrimson within DG (n = 7 mice / 7 sessions). **(D)** Expression of Cre-dependent GCaMP in the LC shown at low magnification (left) and high magnification (right). **(E)** Lack of LC GCaMP response to 20 Hz photic activation of Chrimson within DG (n = 3 mice / 3 sessions; recordings obtained from a subset of animals tested within DG). **(F)** Expression of mCherry (control) and GRAB_NE2m_ in the DG; note the fiberoptic track (arrow). **(G)** Lack of DG GRAB_NE2m_ response to 20 Hz photic stimulation of the mCherry control within DG (n = 4 mice / 4 sessions). For each photic stimulation experiment, displayed are mean (dark lines) and SEM (shaded areas) across all sessions within the group, aligned to stimulation onset **(B1, C1, E1, G1)**, and heatmaps of z_ΔF/F values, in which each row represents data from an individual session **(B2, C2, E2, G2)**. Vertical lines indicate onset of photic stimulation (10 mW, 15 ms pulse width, 10 s).

We next asked whether this locally evoked NE release is mediated by NMDA receptor (NMDAR) activation, as would be consistent with the GANE model. To test this, we again stimulated Chrimson within the DG while recording evoked GRAB_NE2m_ fluorescence, with or without CPP pre-administration (**Fig. 5A**). We observed a significant, ∼50% reduction of fluorescence in the presence of CPP, implicating NMDAR activation in NE release (**Fig. 5B-D**).

**Figure 5.**
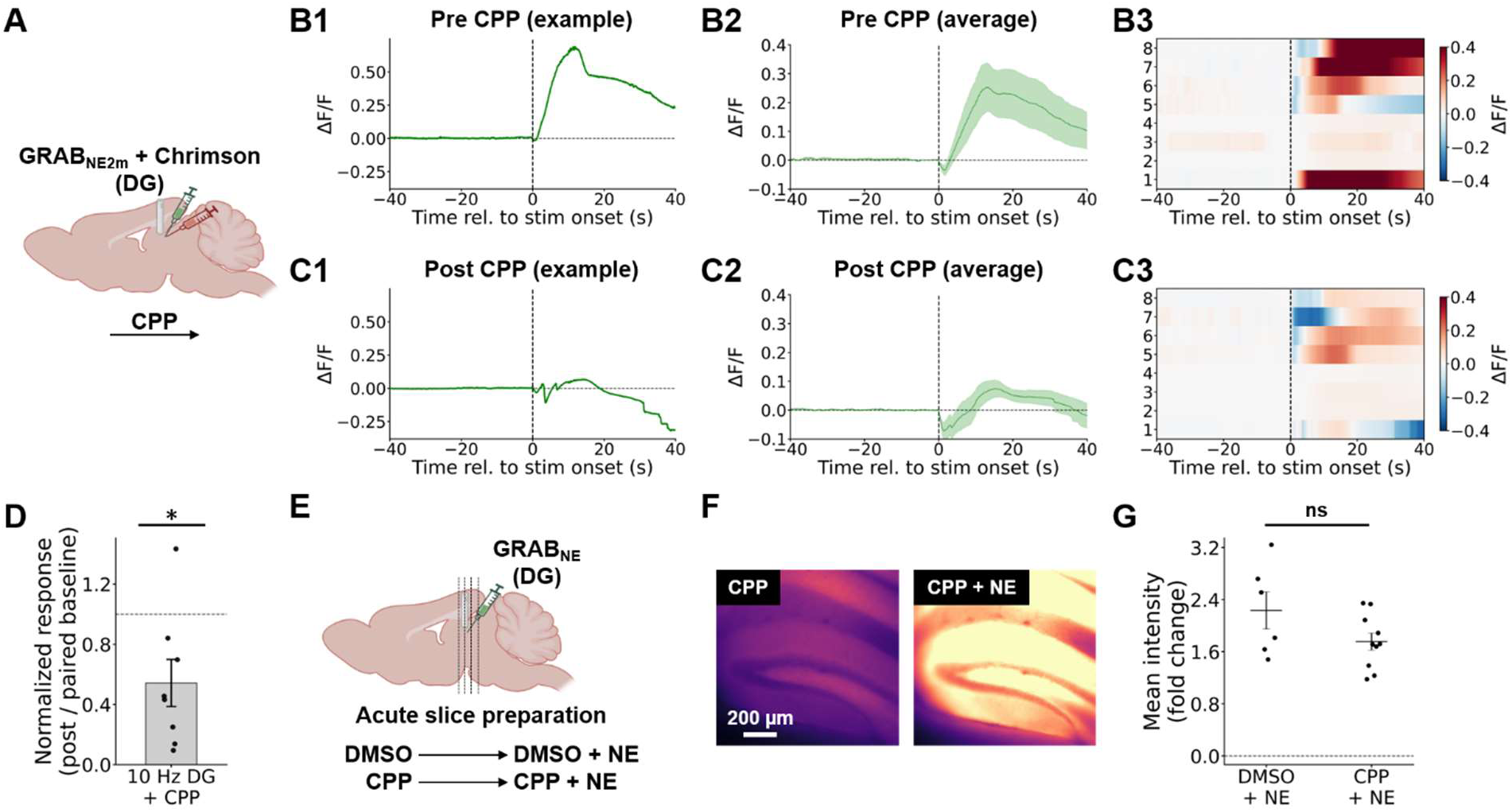
Locally evoked NE release is partially mediated by NMDAR activation. **(A)** Schematic illustrating experimental configuration for local (DG) optogenetic stimulation and concurrent LC-NE circuit recordings, including combined expression of GRAB_NE2m_ and Chrimson in DG and fiberoptic placement above DG (for photic stimulation and fiber photometry). Evoked GRAB_NE2m_ signal in response to photic stimulation within DG delivered under baseline conditions **(B)** or following systemic CPP injection **(C)** (n = 6 mice / 8 sessions). Displayed are individual traces **(B1, C1)**, mean (dark lines) and SEM (shaded areas) across all sessions within the group **(B2, C2)**, and heatmaps of heatmaps of z_ΔF/F values, in which each row represents data from an individual session **(B3, C3)**. Vertical lines indicate onset of photic stimulation (10 Hz, 10 mW, 15 ms pulse width, 10 s). **(D)** Responses in the sessions following systemic CPP injection were normalized by paired baseline responses. CPP injection significantly reduced the evoked response magnitude (p = 0.023, ratio paired t-test). **(E)** Schematic illustrating experimental configuration for acute slice experiments, with injection of GRAB_NE2m_ virus into DG. **(F)** Representative live images showing GRAB_NE2m_ expression in CPP (left) and following NE wash-on (right). **(G)** There was no significant difference in the mean intensity fold-change between (1) vehicle (DMSO) + NE and (2) CPP + NE (p = 0.105, unpaired t-test).

To exclude the unlikely possibility that CPP directly inhibits the GRAB_NE_ sensor, we imaged GRAB_NE2m_ within acute hippocampal brain slices, washing on (1) either vehicle (DMSO; 0.2% in ACSF) or CPP (10 µM), followed by (2) DMSO / CPP plus NE (50µM) (**Fig. 5E**). In slices pre-treated with CPP, NE wash-on caused a robust increase in GRAB_NE2m_ fluorescence (**Fig. 5F**) similar to that observed in the absence of CPP (**Fig. 5G**).

Together, these results support the conclusion that locally evoked NE release is partially mediated by NMDAR activation.

### Ictal NE release occurs across multiple brain regions, epilepsy models, and seizure-evoking methods

If local control of NE release rests upon fundamental neuroanatomical features of the LC-NE circuit, we hypothesized that this ictal phenomenon would generalize across seizure-activated downstream brain regions (within the seizure onset or propagation zone), epilepsy models, and experimental methodology for evoking seizures.

Hyperthermia-evoked seizures in DS mice activate not only the hippocampus, but also multiple neocortical regions, including somatosensory cortex (S1). Using 2P imaging, we found an early increase in GRAB_NE2m_ signal, followed by a dip in fluorescence (**Fig. 6A-C**), as observed in DS DG.

**Figure 6.**
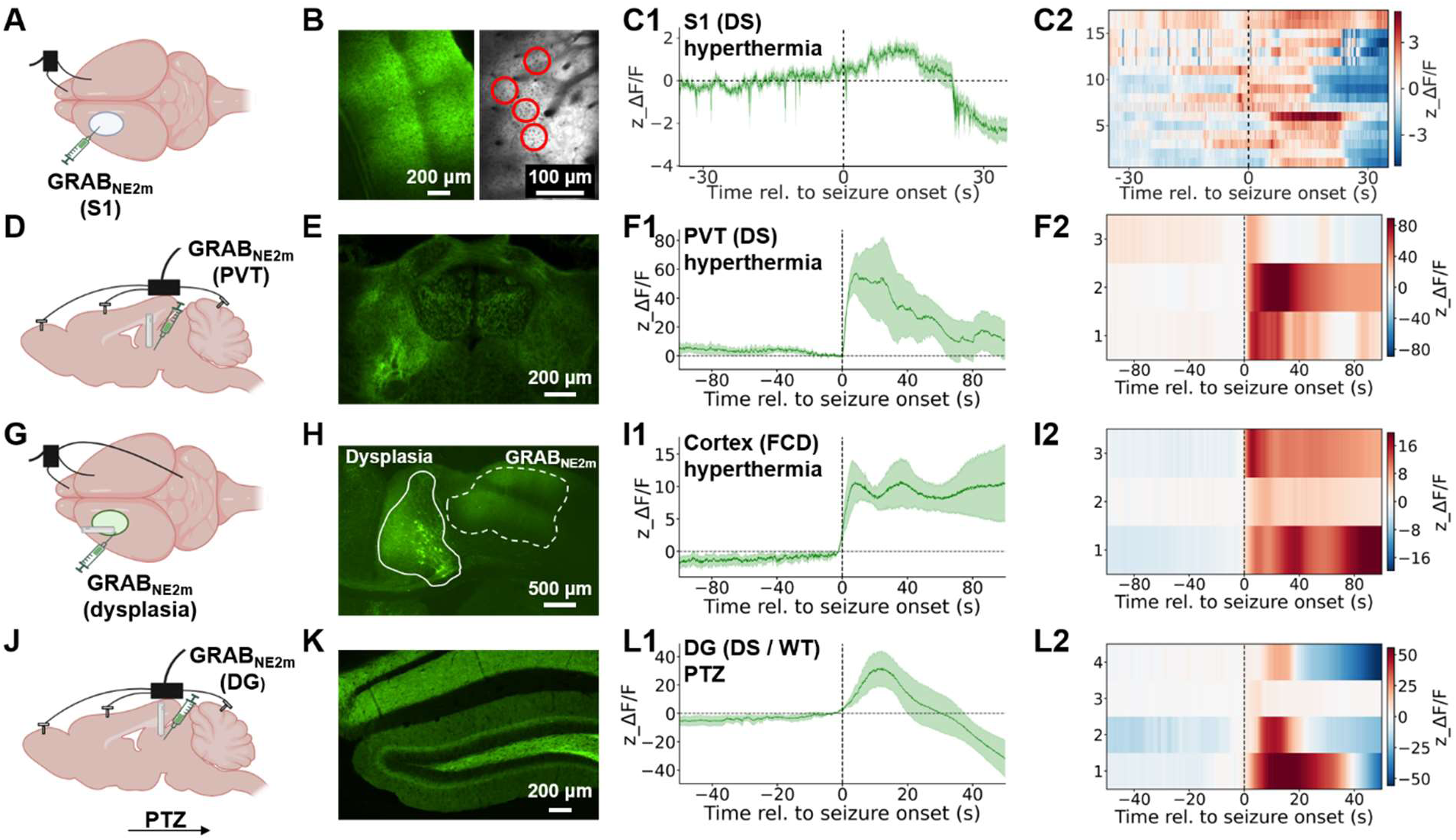
Ictal NE release occurs across downstream brain regions, epilepsy models, and seizure induction methods. **(A)** Schematic illustrating experimental configuration for animals used for 2P recording of GRAB_NE2m_ within wS1, including GRAB_NE2m_ virus injection into wS1, a craniotomy window above wS1, and cortical skull screws for ECoG. **(B)** Expression of GRAB_NE2m_ within wS1, visualized using fluorescence microscopy (left) or 2P imaging in which ROIs are indicated (right). **(C)** Seizure onset-aligned response of ROIs displaying an acute signal increase (n = 17 ROIs / 4 sessions / 2 mice). The AUC of the initial (positive) component of the signal was 20.19 ± 5.36 (p = 0.0008, one-sample t-test). **(D)** Schematic illustrating experimental configuration for NE recordings in PVT in DS mice, including GRAB_NE2m_ virus injection into PVT, fiberoptic placement above PVT, and cortical skull screws for ECoG. **(E)** Expression of GRAB_NE2m_ within PVT. **(F)** Increase in GRAB_NE2m_ signal within PVT in response to hyperthermic seizure induction (n = 3 mice / 3 sessions). **(G)** Schematic illustrating experimental configuration for NE recordings in juxta-dysplastic cerebral cortex in FCD mice, including GRAB_NE2m_ virus injection into cortex, fiberoptic placement above cortex, and cortical skull screws for ECoG. **(H)** Expression of GRAB_NE2m_ adjacent to the dysplasia. **(I)** Increase in GRAB_NE2m_ signal adjacent to the dysplasia in response to hyperthermic seizure induction (n = 3 mice / 3 sessions). **(J)** Schematic illustrating experimental configuration for NE recordings in DG in WT and DS mice, including GRAB_NE2m_ virus injection into DG, fiberoptic placement above DG, and cortical skull screws for ECoG. **(K)** Expression of GRAB_NE2m_ within DG. **(L)** Increase in GRAB_NE2m_ signal within DG in response to PTZ-triggered seizures. As both DS and WT mice exhibited similar results, data were pooled across recordings (n = 2 mice / 2 sessions for WT and 2 DS mice / 2 sessions for DS). For each recording, displayed are mean (dark lines) and SEM (shaded areas) across all sessions within the group, aligned to seizure onset **(C1, F1, I1, L1)**, and heatmaps of z_ΔF/F values, in which each row represents data from an individual session **(C2, F2, I2, L2)**.

The paraventricular thalamic nucleus (PVT), which receives dense inputs from neocortical and limbic regions (Li and Kirouac, 2012), also showed an increase in ictal NE concentrations (**Fig. 6D-F**), demonstrating that this phenomenon occurs within subcortical circuits.

We next sought to investigate ictal NE dynamics in a focal epilepsy model. We employed a mouse model of focal cortical dysplasia, generated via *in utero* electroporation to induce a patient-specific pathogenic mutation (Proietti Onori et al., 2021) resulting in mechanistic target of rapamycin (mTOR) hyperactivation. We recently demonstrated that focal seizures can be acutely triggered via hyperthermia in this model (Zhang et al., 2026). We introduced GRAB_NE2m_ adjacent to the dysplastic region, performed fiber photometry while evoking hyperthermic seizures, and observed a strong associated NE release (**Fig. 6G-I**).

Finally, although our experiments include negative controls for direct temperature effects, it is possible that hyperthermia-evoked seizures differ from other seizure types. To test this, we induced seizures in both DS and WT mice via administration of the chemoconvulsant pentylenetetrazol (PTZ). PTZ-evoked seizures robustly triggered NE release in both genotypes (**Fig. 6J-L)**.

Taken together, these findings demonstrate that ictal NE release is a generalized phenomenon that occurs across multiple downstream brain regions, epilepsies, and evoking measures.

## Discussion

Here, we found that seizures reduced population-level LC-NE GCaMP signal (**Fig. 1**) but triggered supraphysiologic release of NE within downstream regions (**Figs. 2, 6**).

Why might seizures inhibit LC-NE neurons? The relatively slow onset of the GCaMP signal reduction, and its persistence after electrographic seizure offset, suggests several non-mutually exclusive potential mechanisms: recruitment and accumulation of an inhibitory network signal, withdrawal of tonic excitation, a gradually changing chemical state, or an increase in intrinsic autoinhibition. First, seizures may suppress LC-NE activity through direct inhibitory projections. Candidate sources include sleep-promoting GABAergic / galaninergic neurons and melanin-concentrating hormone neurons), the central amygdala or extended amygdala populations, the nucleus prepositus hypoglossi (PrH), and GABAergic neurons of the peri-LC region (Schwarz and Luo, 2015; Chandler et al., 2019; Poe et al., 2020). Second, seizures – or subsequently evoked cortical spreading depression – may suppress cortical and subcortical arousal systems that normally maintain LC firing via direct or polysynaptic projections, including orexin neurons in the lateral hypothalamus, glutamatergic reticular inputs, or prefrontal and limbic inputs. Third, seizures produce extracellular adenosine accumulation, which could suppress LC-NE neuronal activity through direct receptor activation (Shefner and Chiu, 1986), reduced excitatory synaptic input, or effects on astrocytes. Finally, LC-NE neurons are subject to intrinsic autoinhibition mediated by α2-adrenergic autoreceptors (e.g., (Arima et al., 1998)) such that any seizure-driven NE release within LC would further inhibit LC-NE activity. Future studies will be needed to determine the relative contributions and temporal dynamics of these different candidate mechanisms.

An important limitation of our approach is that fiber photometry provides a bulk signal, so our data may obscure heterogeneity of response to seizures (as was recently observed in a temporal lobe epilepsy model (Larsen et al., 2023)). Indeed, LC-NE neurons comprise distinct subpopulations that differ in their projection targets and behavioral functions (Schwarz and Luo, 2015; Chandler et al., 2019; Poe et al., 2020; Su et al., 2026). Future studies restricting GCaMP expression to projection-based or functionally specialized subpopulations will be necessary to further resolve and understand seizure-related dynamics within the LC.

Reconciling the dissociation between somatic inhibition and terminal signals is central to understanding LC-NE circuit function during seizures. Fluorescent reporters of NE (and other neuromodulators) enable high temporal resolution readout *in vivo*, but – as our work illustrates – require multiple controls to ensure interpretability of results. It is now broadly accepted that isosbestic recordings are essential to correct for motion artifacts (especially relevant for convulsive seizures), photobleaching, autofluorescence, and other artifactual sources of signal variability. However, ictal recordings require additional scrutiny. Seizures introduce pH and hemodynamic shifts beyond the physiologic range in which fluorescent sensors have been optimized and validated (Zhang et al., 2022; Byron and Sakata, 2024; Zhuo et al., 2024), and the resulting optical artifacts may not fully rectified by isosbestic correction (**Fig. 3; Supplementary Fig. 2**). Similar event-onset associated dips in fluorescent signal have been recently reported during seizures, with exogenous stimulation, and even following changes in physiologic state (Wu et al., 2023; Zhuo et al., 2024; Huang et al., 2026; Neyhart et al., 2026). Red-shifted sensors, such as the newly developed GRAB_rNE2.0_ (**Fig. 3**), may be less vulnerable to these sources of artifact (**Fig. 3**).

The supraphysiologic magnitude of the ictal NE release (**Fig. 2**) argues that it is not simply derived from a subpopulation of LC-NE neurons, the activation of which was not detected (**Fig. 1; Supplementary Fig. 5**), or from a noradrenergic source beyond the LC or NTS (**Supplementary Fig. 4**). A more parsimonious explanation is that ictal activity within downstream regions locally evokes NE release, a concept in fact experimentally demonstrated over 40 years ago when ictal NE release was evidenced even after acute transection of the dorsal noradrenergic bundle (Nelson et al., 1980).

The GANE model posits a physiological effect of glutamate-induced local NE release for information processing, and seizures may overwhelm this mechanism due to supraphysiologic ictal glutamate release. *In vivo* support for the GANE model includes prior work demonstrating that visual stimulation enhanced vigilance-dependent NE release within primary visual cortex (Gray et al., 2021) and that increased hippocampal spiking amplified NE signaling evoked by contextual arousal (Huang et al., 2026). It is notable that high frequency (20 Hz) photoactivation in our experimental preparation was sufficient to evoke NE release, but that low frequency (1 Hz) activation was not (**Fig. 4**). Future experiments will be needed to more fully explore the stimulation parameter space, and to establish conditions – such as presence or absence of evoked after discharges – required to evoke NE release.

The GANE model is premised upon foundational studies demonstrating that glutamate receptor agonists triggered NE release within acute slice and synaptosomal preparations in which the LC was anatomically absent (Gray and Whittaker, 1962; Vezzani et al., 1987; Fink et al., 1989, 1992; Pittaluga and Raiteri, 1990, 1992; Göthert and Fink, 1991; Wang et al., 1992; Andrés et al., 1993; Pittaluga et al., 1999; Luccini et al., 2007). Our finding that NE release in DG was partially inhibited by NMDAR antagonism (**Fig. 5**) is consistent with this early work, and with a more recent study demonstrating that local NMDAR antagonism reduced visual stimulation-evoked NE release in visual cortex (Gray et al., 2021). We therefore hypothesize that ictal NE release is a widespread phenomenon and will follow ictal glutamate release wherever it occurs across brain regions and seizure types. Indeed, we demonstrated seizure-evoked NE release across multiple experimental conditions: within DG (**Fig. 2**), S1 (**Fig. 6A-C**), and PVT (**Fig. 6D-F**) of DS mice with hyperthermic seizures, the juxta-dysplastic cortex of FCD model mice with hyperthermic seizures (**Fig. G-I**), and the DG of both DS and WT mice with PTZ-evoked seizure (**Fig. 6J-L**). Future studies will be needed to more thoroughly characterize NE release dynamics in these different conditions, noting potential interesting heterogeneity in signal amplitude and kinetics (**Fig. 6**). It will also be informative to assay NE levels in brain regions that are not directly activated by seizures, or in seizure models (e.g., (Meyer et al., 2018)) that involve hypersynchrony but not hyperactivity.

Notably, there are multiple additional mechanisms by which seizures may locally activate terminals independently of somatic spikes. Local spikes may be generated ectopically within distal axons. Axons or boutons may be directly depolarized through extreme elevation of extracellular potassium (e.g., (Duan et al., 2026)). Cytosolic NE may leave terminals through reverse transport through the norepinephrine transporter (NET), as occurs with amphetamine (Robertson et al., 2009). Spreading depolarization may co-occur with seizures – as previously observed in DS mice (Aiba et al., 2023) – and indiscriminate trigger neurotransmitter release via depolarization of terminals. Future studies will be necessary to examine the relative contributions of these different mechanisms.

What are the expected functional consequences of seizure-evoked somatic LC-NE inhibition and supraphysiologic NE release? Inhibition of LC-NE neurons plays a crucial role in physiologic sleep (e.g., (Carter et al., 2010) and loss of consciousness from general anesthesia (Du et al., 2018), so it stands to reason that ictal inhibition of LC-NE neurons would likewise suppress arousal. High-frequency excitatory stimulation of LC-NE neurons paradoxically caused a behavioral arrest, likely due to depletion of NE stores (Carter et al., 2010), so supraphysiologic ictal release of NE may also impair consciousness until baseline circuit function returns. To confirm these hypotheses, it will be important to examine LC-NE circuit dynamics in seizures with preserved consciousness, to track postictal recovery of circuit function with recovery of consciousness, and to test the impact of pharmacologic or other manipulations of LC-NE signaling. Future studies will also be needed to clarify the peri-ictal dynamics of other neuromodulators that contribute to the ascending arousal system; as presynaptic glutamate receptors also modulate serotonergic and dopaminergic neurotransmission (Ohta et al., 1994; Meltzer et al., 1997; Martín-Ruiz et al., 2001; de Kock et al., 2006), we hypothesize that the same pattern will be seen for ictal serotonin and dopamine release. Such work will inform the development of strategies to preserve arousal both during and after seizures.

## Acknowledgements

We are grateful to Dr. Yu Wang for providing the FCD model mice used in this manuscript, and to the entire Mattis laboratory for helpful discussions.

## CRediT author statement

**Meiling Zhao:** conceptualization, methodology, software, validation, formal analysis, investigation, data curation, writing – original draft, visualization. **Yiannos Demetriou:** software, formal analysis, data curation, writing – original draft, visualization. **Anam Noor:** validation, investigation, writing – review and editing. **Joseph Barden:** validation, formal analysis, investigation, visualization, writing – review and editing. **Limei Zhu:** investigation, visualization, writing – review and editing. **Qianyi Yu:** formal analysis, investigation, writing – review and editing. **Alexander Sajdak:** software, formal analysis, writing – review and editing. **Arnav Akhaury:** investigation, visualization, writing – review and editing. **Chandni Rana:** methodology, software, writing – review and editing. **Peilin Yang:** resources, writing – review and editing. **Yulong Li:** resources, writing – review and editing. **Christian Burgess:** conceptualization, methodology, writing – review and editing. **Sam McKenzie:** conceptualization, methodology, writing – review and editing. **Paul Kramer:** conceptualization, methodology, validation, formal analysis, investigation, data curation, writing – original draft, visualization. **Joanna Mattis:** conceptualization, methodology, formal analysis, resources, writing – original draft, visualization, supervision, project administration, funding acquisition.

