## Supplemental materials for "Seizures in a Dravet Syndrome mouse model inhibit the locus coeruleus but locally evoke supraphysiologic norepinephrine release"

### Supplemental figures and tables

#### Supplemental Figure 1. Example LC-NE GCaMP fiber photometry signals, ECoG traces, and temperature curves.

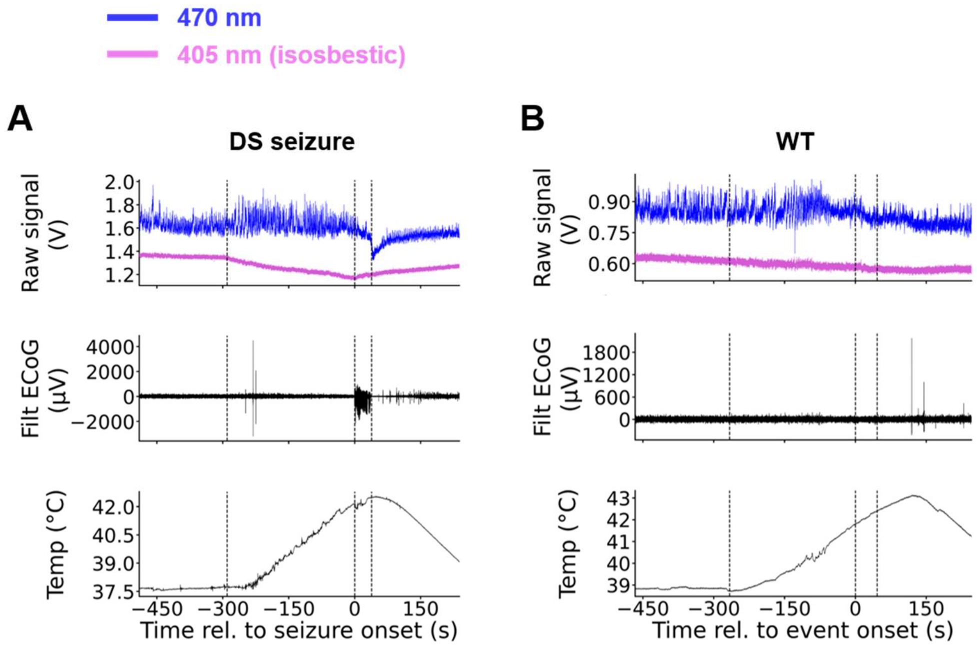


**(A)** Representative GCaMP recording from a DS mouse that exhibited a hyperthermic seizure. **(B)** Representative recording from a WT control mouse that was heated but did not exhibit a seizure. Synchronized data include (1) 470 nm and 405 nm (isosbestic) signals recorded from GCaMP expressed within LC-NE neurons (top), (2) ECoG (middle), and (3) temperature. The left-most vertical dashed line indicates heating onset; the second and third vertical dashed lines indicate seizure onset and offset or equivalent temperatures.

Supplemental Figure 2. Interictal activity and intrinsic excitability of LC-NE neurons is normal in DS mice.

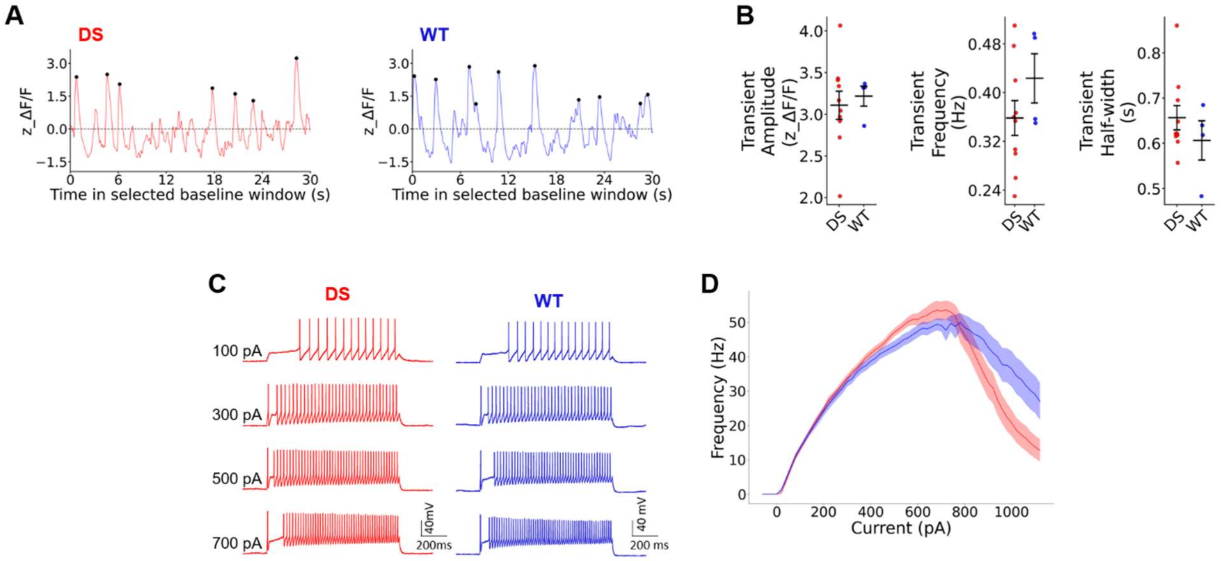


**(A)** Representative interictal GCaMP transients recorded from LC-NE neurons in DS mice (left) and WT mice (right) from DG *in vivo*, illustrating no evident genotype differences. **(B)** Interictal GCaMP amplitude, frequency, and half-width were similar between DS and WT LC-NE neurons. n = 8 mice / 10 sessions (DS) and 3 mice / 4 sessions (WT). **(C)** Representative current clamp responses to depolarizing current injections in DS and WT LC-NE neurons. **(D)** Current/frequency plots were similar between DS and WT LC-NE neurons (genotype effect p = 0.29; mixed effects model).

Supplemental Figure 3. Seizures trigger an artifactual signal reduction in $GRAB_{NE}$.

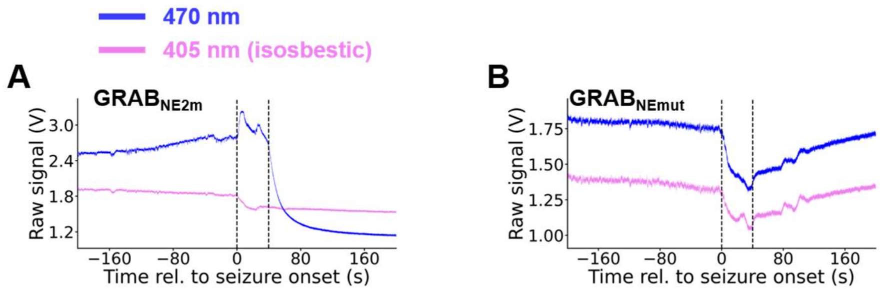


**(A)** Representative raw 470 nm and 405 nm signals recorded from $GRAB_{NE2m}$ in DG. A transient signal reduction was observed in the 405 nm (isosbestic) signal following seizure onset. **(B)** Representative raw 470 nm and 405 nm signals recorded from $GRAB_{NEmut,}$ in DG. A signal reduction was observed in both raw 405nm and 470nm traces following seizure onset.

Supplemental Figure 4. Absence of evidence for direct bilateral projections between DG and NTS-NE neurons.

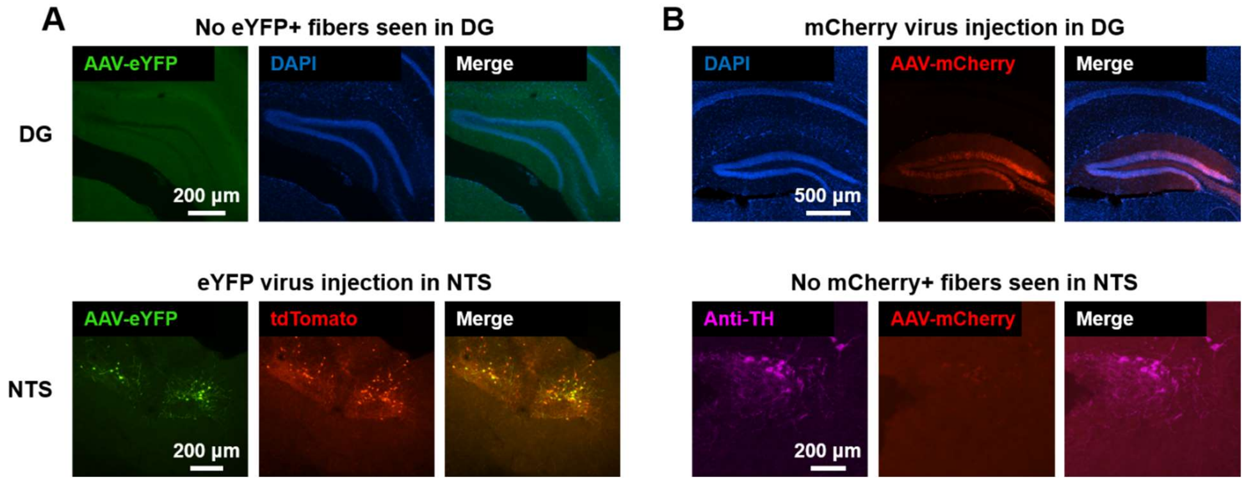


**(A)** Cre-dependent eYFP-expressing virus was injected into the NTS of a *Dbh*-Cre;tdT mouse. Within NTS (below), there was overlap in expression between YFP (virus) and tdT (Dbh marker), as expected. Within DG (above, outlined anatomically by DAPI, blue), no eYFP expression was observed. **(B)** An mCherry-expressing virus (red) was injected into the DG of a WT mouse DG (outlined anatomically by DAPI, blue). No mCherry expression was observed within the NTS (identified by immunostaining for TH, purple).

Supplemental Figure 5. Hyperthermic seizures do not induce elevated c-Fos expression in LC-NE neurons.

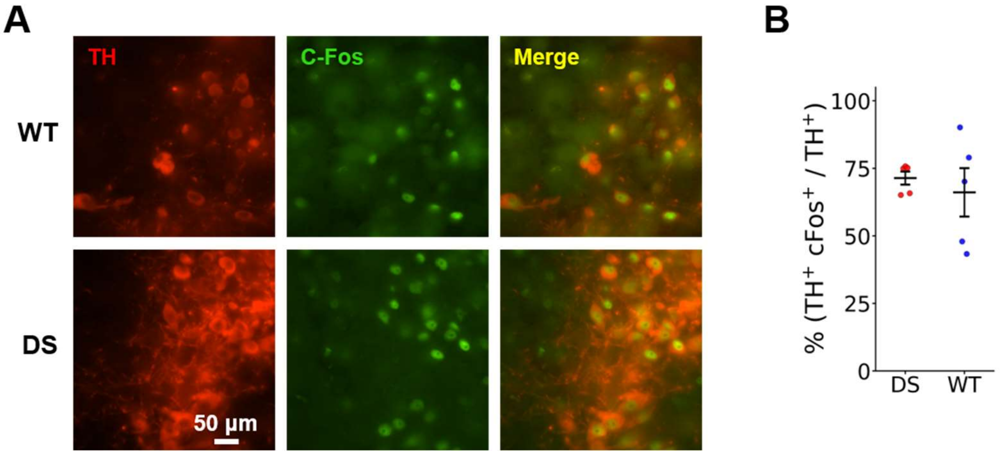


**(A)** High-magnification (60x) images showing immunohistochemical labeling of LC-NE neurons via tyrosine hydroxylase (TH, red) and the subset of recently active neurons using the transcription factor c-Fos (green) as a proxy for neuronal activity. **(B)** Quantification of the proportion of LC-NE ($TH^{+}$) neurons expressing c-Fos in n = 5 DS mice and n = 5 non-seizing WT littermates following a single hyperthermic seizure induction session. An unpaired t-test found no significant difference between DS and WT mice with respect to the proportion of LC-NE neurons expressing c-Fos ($p = 0.5840$).

Supplemental Table 1. Intrinsic physiologic properties of LC-NE neurons in DS and WT mice.

| Measurement | DS | WT | Corrected p-value |
|---|---|---|---|
| n (mice / cells) | 5 / 19 | 5 / 20 | |
| **Intrinsic Property** | | | |
| Cm (pF) | 46.25 ± 1.31 | 47.49 ± 2.16 | 0.94 |
| Rheobase (pA) | 39.68 ± 3.66 | 35.34 ± 3.45 | 0.94 |
| Rm (MOhm) | 749.09 ± 54.97 | 778.34 ± 85.01 | 0.94 |
| Sag | 0.99 ± 0.002 | 0.99 ± 0.002 | 0.94 |
| Time Constant (ms) | 46.48 ± 1.88 | 47.88 ± 2.01 | 0.94 |
| Vm (mV) | -62.37 ± 1.94 | -60.87 ± 1.70 | 0.94 |
| **Repetitive AP Property** | | | |
| ISI_CoV | 0.19 ± 0.05 | 0.23 ± 0.05 | 0.95 |
| Initial Burst Length (ms) | 365.73 ± 109.36 | 363.71 ± 111.14 | 0.99 |
| Max Instantaneous (Hz) | 89.58 ± 3.02 | 85.94 ± 4.19 | 0.95 |
| Max Steady-state (Hz) | 62.00 ± 1.60 | 60.37 ± 2.06 | 0.95 |
| Maximal Burst Length (ms) | 940.07 ± 9.38 | 943.06 ± 8.14 | 0.95 |
| SFA10 | 2.02 ± 0.43 | 2.55 ± 0.48 | 0.95 |
| SFAn | 2.31 ± 0.52 | 2.61 ± 0.50 | 0.95 |
| **Individual AP Property** | | | |
| AHP Amplitude (mV) | 10.13 ± 0.44 | 9.34 ± 0.73 | 0.55 |
| AP Amplitude (mV) | 66.51 ± 1.10 | 69.30 ± 1.12 | 0.25 |
| AP Half-Width (ms) | 1.21 ± 0.03 | 1.13 ± 0.03 | 0.25 |
| AP Peak (mV) | 37.81 ± 0.77 | 41.47 ± 0.83 | 0.25 |
| AP Rise Time (ms) | 0.91 ± 0.02 | 0.84 ± 0.02 | 0.25 |
| AP Threshold (mV) | -28.70 ± 0.82 | -27.83 ± 0.86 | 0.53 |
| APD 50 (ms) | 1.68 ± 0.03 | 1.57 ± 0.03 | 0.25 |
| APD 90 (ms) | 2.17 ± 0.03 | 2.05 ± 0.04 | 0.25 |

Intrinsic properties, properties of repetitive APs, and individual AP properties calculated for LC-NE neurons from DS (n = 5 mice / 19 cells) and WT (n = 5 mice / 20 cells). No significant differences were observed across any measure.
